# Wnt11 Positively Regulates Neonatal Cardiomyocyte Maturation at the Interphase of Life via Frizzled 4 Receptor

**DOI:** 10.1101/2025.06.01.657323

**Authors:** Xuedong Kang, Joan Moci, Charlotte Wolf, Marlin Touma

## Abstract

Congenital heart defects (CHDs) affect 1% of live births and remain the leading cause of infant morbidity and early mortality. While most studies focus on the genetic basis of CHDs, relatively little is known about the interplay between intrinsic signaling and external environmental factors in the progression of CHDs after birth during the perinatal circulatory transition window when environmental stress factors are prevalent. We recently explored such interplay through a newly identified gene-environment regulatory circuit involving Wnt11 signaling and systemic hypoxia. Specifically, we demonstrated that activation of the Wnt11/Rb1 axis is critical for normal chamber-specific development after birth. This regulatory switch is disrupted by systemic hypoxia more robustly in the right ventricle (RV) than the left ventricle (LV), leading to enhanced neonatal cardiomyocyte cell cycle activity in an RV-specific manner, resulting in delayed maturation and attenuation of ventricular patterning in response to systemic hypoxia stress in the neonatal heart. Furthermore, we found that the Wnt11/Rb1 axis is also inactivated in infantile hearts with cyanotic CHDs, such as tetralogy of Fallot (TOF), potentially contributing to hypoxia-associated RV abnormalities in this context. However, the molecular players of this signaling cascade in neonatal cardiomyocyte remain largely unknown. Herein, we report that Frizzled 4 (Fzd4) acts as a specific upstream receptor for Wnt11 in neonatal cardiomyocytes. Specifically, Fzd4 exhibited an expression pattern like Wnt11 in neonatal heart perinatal circulatory transition under normal and hypoxemic environments. Furthermore, Fzd4 loss in neonatal cardiomyocytes stimulated cardiomyocyte cell cycle activity and disrupted the Wnt11-Rb1 signaling axis mirroring the impact of the Wnt11-deficient cardiomyocyte phenotype. Finally, co-immunoprecipitation analysis confirmed the Wnt11-Fzd4 binding in isolated neonatal cardiomyocytes and intact hearts. These results demonstrate that Fzd4 is a specific and required upstream receptor for the Wnt11-Rb1 signaling activity in the neonatal heart and provides mechanistic insights into the essential role of Wnt11 as a key positive regulator of neonatal cardiomyocyte transition from proliferative to mature phenotype at the interphase of life.

## Introduction

Soon after birth, the newborn heart encounters rapid changes in hemodynamic load and external environment [1-4]. In tandem, elaborate changes in neonatal cardiomyocytes affect proliferative capacity, morphology, cellular metabolism, and ion handling, leading to binucleation, cell cycle exit, and terminal differentiation. These changes are different in temporal pattern and magnitude between the RV and LV [1-8]. However, the underlying regulatory pathways and their potential roles in the pathogenesis of CHDs during perinatal heart chamber development have not been established.

In previous work [1-3], Wnt11, a main player of the non-canonical, β-catenin independent, Wnt pathway [9-13], was identified as an important regulator of chamber-specific perinatal heart growth and development, acting as a negative regulator of the neonatal cardiomyocyte cell cycle genes via enhancing Rb1 activity (by dephosphorylation) more robustly in the neonatal RV chamber [1-3]. In contrast, a perinatal hypoxic environment suppressed the Wnt11-Rb1 axis and induced cardiomyocyte cell cycle activity in an RV-specific manner in the neonatal heart. Importantly, we demonstrated that the Wnt11-Rb1 axis is also suppressed in the hypoxemic human outflow tract specimens obtained from infants with cyanotic tetralogy of Fallot (TOF), indicating potential translational relevance for this new Wnt11-hypoxia regulatory circuit in cyanotic CHDs. Indeed, previous studies have shown that Wnt11 inactivation in mice leads to outflow tract abnormalities and conotruncal defects, including pulmonary stenosis, double outlet right ventricle (DORV), and TOF with early lethality [13]. However, the role of Wnt11 in hypoxemia-induced RV abnormalities in cyanotic TOF cases after birth is entirely unknown. Our findings suggest that the Wnt11 signaling cascade is regulated by hypoxia, and it is plausible that several key molecular players in this cascade remain unknown, and their functional roles in neonatal cardiomyocytes during the perinatal maturation are yet to be identified.

Here, we set out to elucidate the role of the 7-pass transmembrane Frizzled 4 (Fzd4) protein, a member of the G protein-coupled receptors (GPCRs) superfamily [14], as a key upstream receptor for Wnt11 signaling-mediated regulation of cardiomyocyte cell cycle activity during perinatal heart maturation.

## Results

### Fzd4 as a potential receptor for Wnt11 in neonatal cardiomyocytes

Wnt ligands exert their signaling activity via the frizzled (Fzd) receptors, of which Fzd2 and Fzd7 have been identified as noncanonical Wnt receptors [15,16]. However, among eight known Fzd receptors that we screened, *Fzd4* exhibited the highest expression in neonatal mouse RVs at postnatal day 3 (P3) [Figure 1.A, B], like Wnt11 [Figure 1. C].

**Figure 1.**
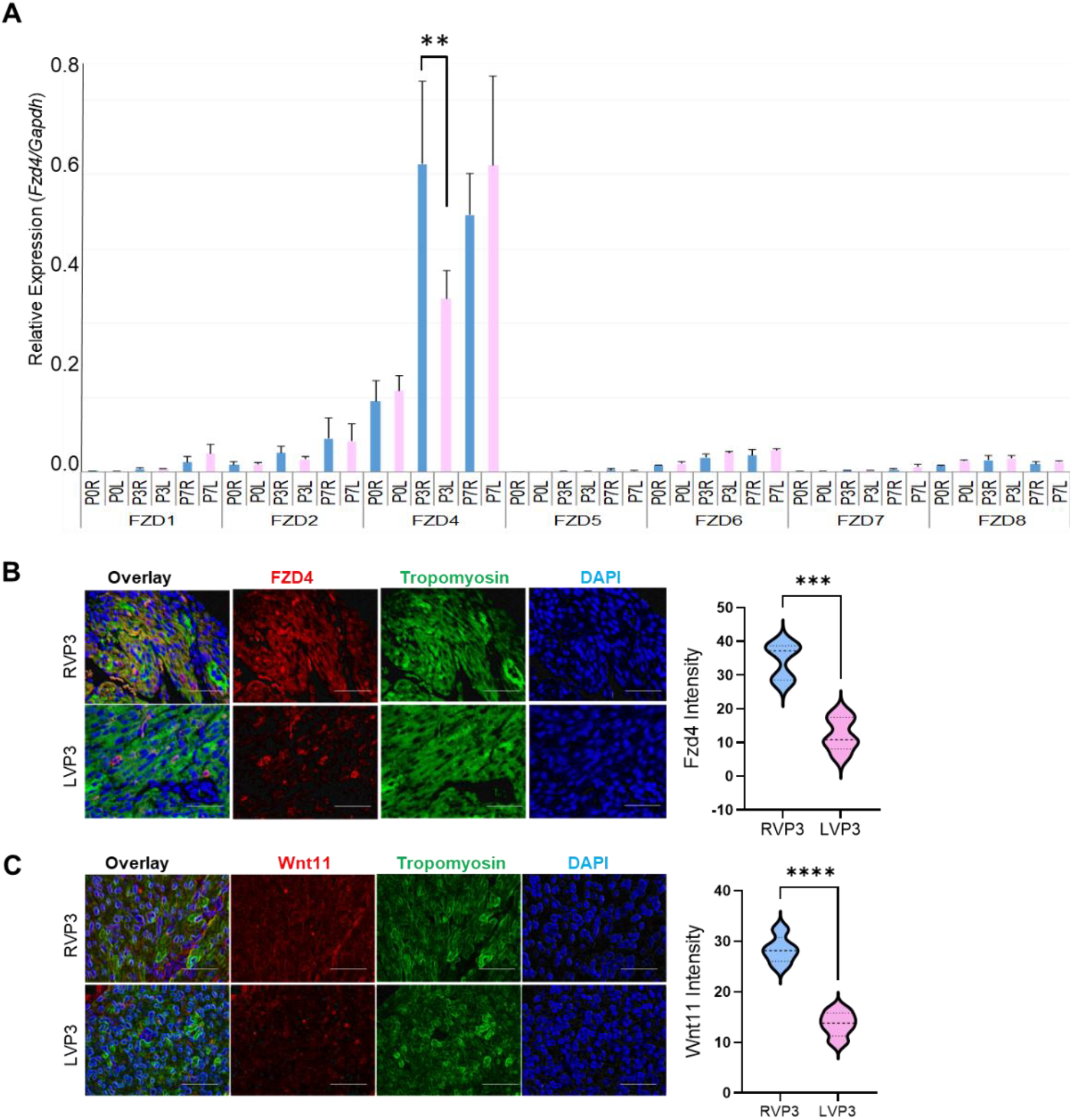
Chamber-specific enrichment of Fzd4 and Wnt11 in neonatal right ventricle (RV) at Postnatal day 3 (P3). (**A**) qRT-PCR analysis of *Fzd4* in LV or RV myocardium of WT neonatal mouse at P0, P3, and P7. Blue, RV; pink, LV. Error bars represent SD. (**B**) Representative confocal images of anti–Fzd4 immunohistochemistry (IHC) and Fzd4 intensity analysis in RVs or LVs of WT neonatal mouse heart at P3. Original magnification: ×40. (**C**) Representative confocal images of anti–Wnt11 IHC and Wnt11 intensity analysis in RVs or LVs of WT neonatal mouse heart at P3. Original magnification: ×40. *n* = 3 replicates per condition. \*\**P* ≤ 0.01; \*\*\**P* ≤ 0.005; \*\*\*\**P* ≤ 0.0001.

### Fzd4 expression is suppressed in response to perinatal systemic hypoxia

Perinatal systemic hypoxia [FIO2:10%] increased Hif1a stabilization in neonatal hearts at P3 [Figure 2.A] and suppressed Wnt11 and induced cardiomyocyte proliferation more robustly in the RV than in the LV [Figure 2. B].

**Figure 2.**
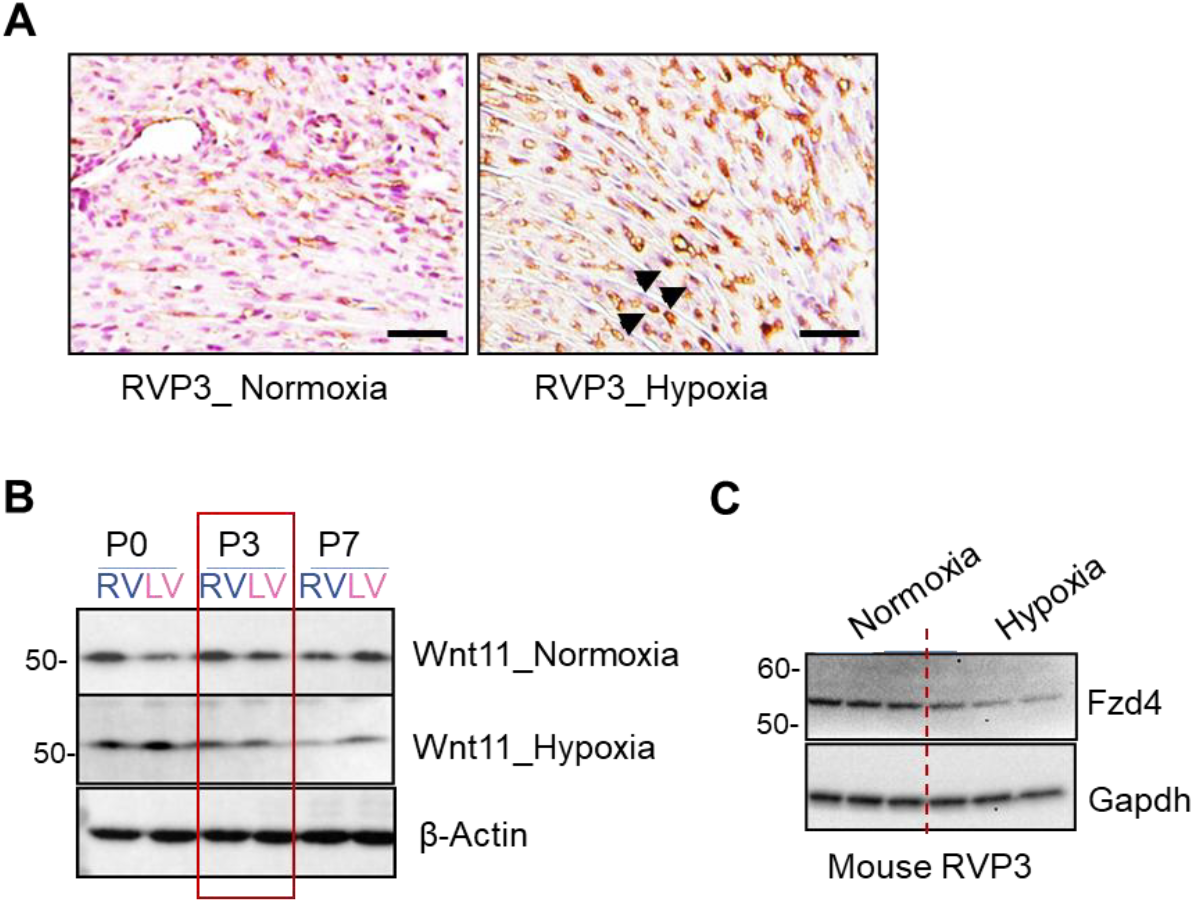
Perinatal systemic hypoxia suppresses Fzd4 and Wnt11 expressions in neonatal right ventricle at P3. (**A**) Representative light microscopy images of anti–Hif1α immunohistochemistry (IHC) in RVs of neonatal mouse at P3 in normoxic and hypoxic conditions. Arrowheads indicate Hif1α-positive cardiomyocytes. Original magnification, ×40. (**B**) Expression of Wnt11 protein (Western blot) in LV and RV myocardium of WT neonatal mouse hearts at P0, P3, and P7 (Western blot). *n* = 3 replicates per condition. Adopted from reference [1]. (**C**) Expression of Fzd4 protein (Western blot) in RV myocardium of WT neonatal mouse at P3 in normoxic and hypoxic conditions. (Western blot). β-Actin or Gapdh were used as loading control. *n* = 3 replicates per condition.

Remarkably, like Wnt11, Fzd4 protein expression was also suppressed more robustly in the RV at P3 in response to perinatal systemic hypoxia [Figure 2. C].

### Fzd4 binds to Wnt11 in neonatal cardiomyocytes

Fzd4 receptor has been linked to the noncanonical Wnt11/PCP signaling during vascular development [17]. Whether Wnt11 interacts with the Fzd4 receptor in neonatal cardiomyocytes is unknown. Co-immunoprecipitation (co-IP) assays confirmed that Fzd4 protein directly binds to Wnt11 in cellular lysates from isolated neonatal cardiomyocytes [Figure 3.A] as well as neonatal mouse RV at P3 [Figure 3.B]. Thus, Fzd4 is a candidate receptor for Wnt11 signaling in neonatal cardiomyocytes.

**Figure 3.**
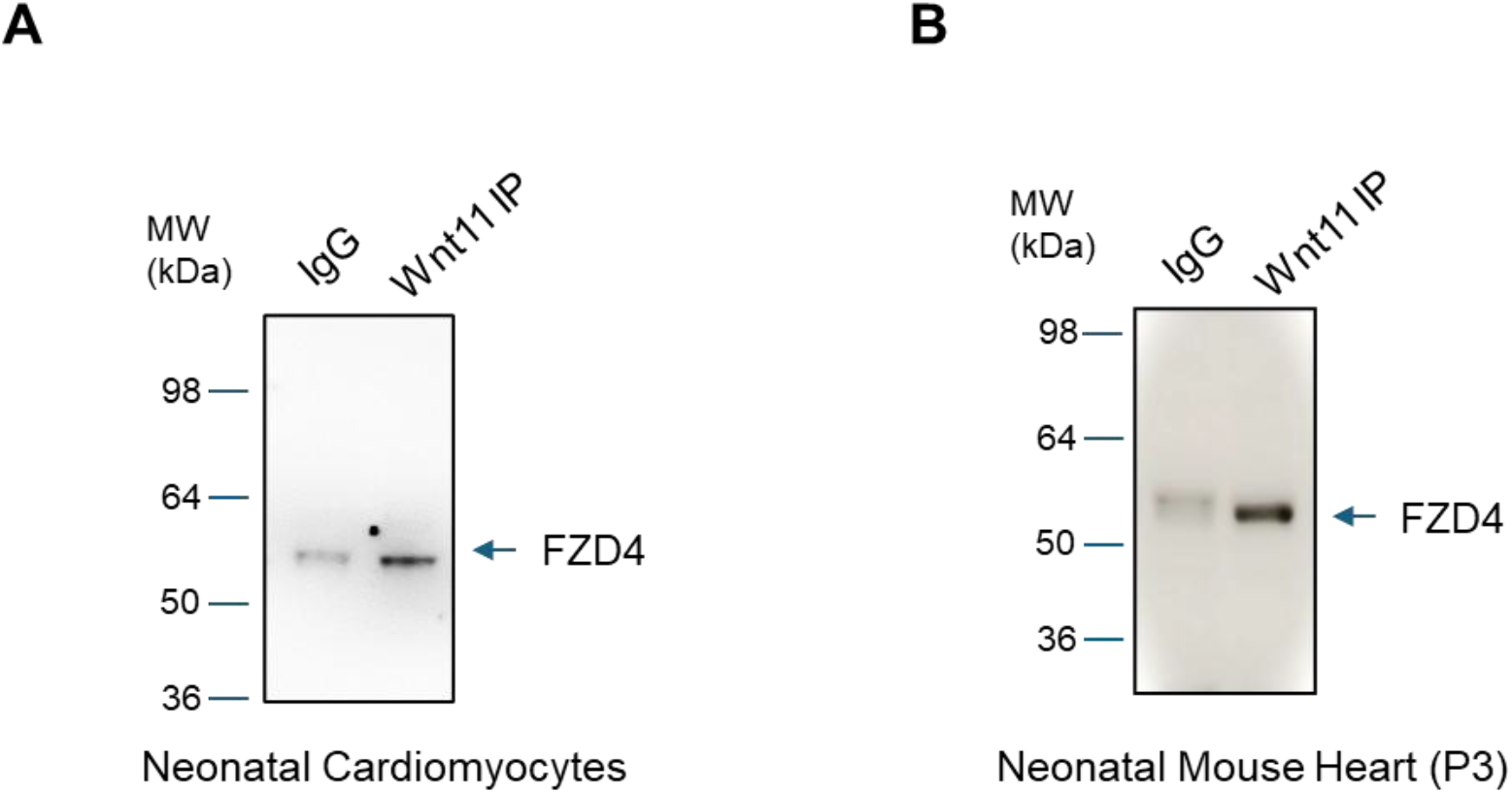
Co-immunoprecipitation (co-IP) analysis confirm Wnt11-Fzd4 binding in neonatal heart. (**A**) co-IP analysis depicts the presence of Fzd4 protein in the Wnt11-precipitated material recovered from neonatal cardiomyocyte lysate, was assessed by western blotting with anti-FZD4 antibody. (**B**) co-IP analysis depicts the presence of Fzd4 protein in the Wnt11-precipitated material recovered from neonatal mouse cellular lysate, was assessed by western blotting with anti-FZD4 antibody.

### Fzd4 loss in neonatal cardiomyocytes induces cell cycle activity

We have previously shown that Wnt11 inhibition in neonatal cardiomyocytes and in neonatal hearts induces cardiomyocyte cell cycle activity [1]. To test whether Fzd4 loss is sufficient to induce cardiomyocyte cell cycle activity, we suppressed *Fzd4* in primary cultured neonatal cardiomyocytes using *Fzd4*-specific siRNA. Like the phenotype observed in the Wnt11-deficient neonatal cardiomyocytes, Fzd4 loss induced cardiomyocyte cell cycle activity as demonstrated by increased expression of the mitosis marker Aurora Kinase B (AurkB) and increased AurkB-positive cardiomyocytes indicating cell cycle activation [Figure 4. A, B]. To determine if Fzd4 is required for Wnt11-mediated regulation of neonatal cardiomyocyte cell cycle activity, we examined whether exogenous recombinant Wnt11 (rWnt11) may mitigate the impact of Fzd4 loss. The results indicated that rWnt11 treatment failed to rescue the Fzd4-deficient phenotype [Figure 4.A], supporting that the Fzd4 receptor is specifically required for Wnt11 signaling in neonatal cardiomyocytes.

**Figure 4.**
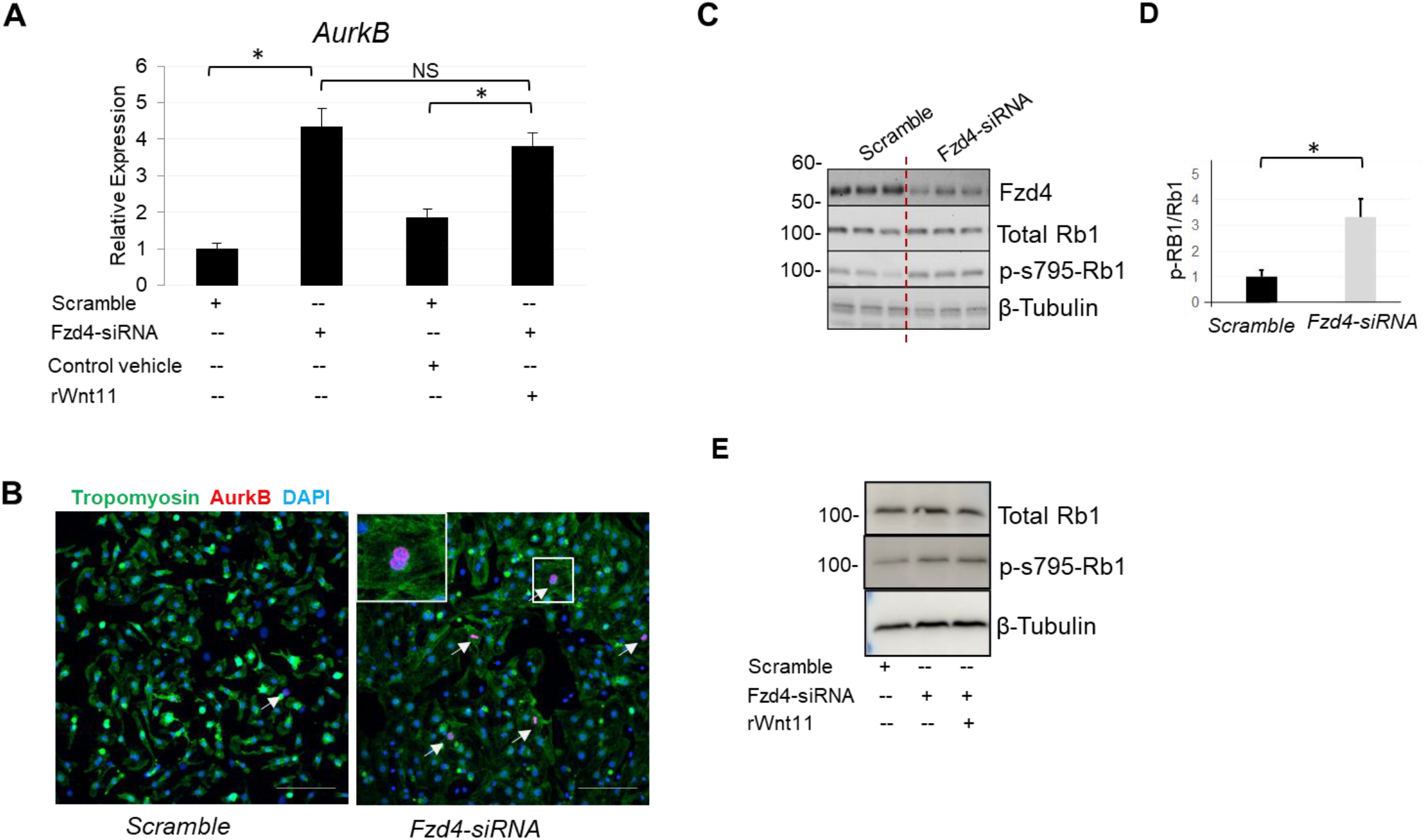
Fzd4 inhibition induces neonatal cardiomyocyte proliferation and disrupts the Wny11-Rb1 signaling axis. (**A**) Expression analysis of the mitosis marker (AurkB) using RNA from scramble-treated or *Fzd-siRNA*–treated NRVMs with/without exogenous rWnt11 treatment (qRT-PCR). (**B**) Representative confocal images of AurkB–positive cardiomyocytes (arrows) in scramble-treated or Wnt11 siRNA–treated NRVMs. (**C**) Expression of Fzd4, Rb1 protein, and phosphorylated Rb1 (p-Rb1) at Ser795, using protein from scramble-treated or *Fzd-siRNA*–treated NRVMs (Western blot). (**D**) quantification analysis of p-Rb1/Rb1 ratio in scramble-treated or Wnt11 siRNA–treated NRVMs. (**E**) Expression of Rb1 protein and p-Rb1 at Ser795, using protein from scramble-treated or *Fzd-siRNA*–treated NRVMs with/without exogenous rWnt11 treatment (Western blot). β-Tubulin was used as loading control. Original magnification, x40. N= 3 replicates per condition. \**P* ≤ 0.05.

### Fzd4 loss disrupts the Wnt11-Rb1 axis

Wnt11 loss in neonatal cardiomyocytes promotes the cell cycle via increasing phosphorylated Rb1(p-RB1), which is an inactive form of Rb1. In the differentiated, mature, cardiomyocytes, Rb1 binds to the transcription factor E2f1 and suppresses its transcriptional activity. When Rb1 is phosphorylated, it separates from E2f1, which in turn translocate to the nucleus and activates the cell cycle progression program [1]. To determine the functional impact of Fzd4 loss on the Wnt11-Rb1 signaling axis, we examined whether Fzd4 loss may alter Rb1 expression and/or phosphorylation state. We selected the phosphorylation site s795, which was previously shown to be required for E2f1-spicific binding [18,19] and blocked by Wnt11 manipulation in neonatal cardiomyocytes [1]. While total Rb1 protein was not affected, all Rb1 phosphorylated (inactive) forms were significantly increased [Figure 4. C, D]. Furthermore, exogenous rWnt11 treatment failed to rescue Rb1 activity in the Fz4-deficient neonatal cardiomyocytes [Figure 4. E]. Together, these findings indicate that Fzd4 loss disrupts the Wnt11-Rb1 axis in neonatal cardiomyocytes. Therefore, Wnt11 exerts signaling activity in neonatal cardiomyocytes via the Fzd4 receptor. The Fzd4 receptor is a specific, required, and nonredundant receptor for Wnt11 signaling regulation of neonatal cardiomyocyte cell cycle exit.

## Discussion

In this study, we identify Fzd4 as a specific, required, and nonredundant Wnt11 receptor in neonatal cardiomyocytes. First, the Fzd4 expression pattern in the neonatal heart mirrors Wnt11 under normoxic and hypoxic conditions. Second, Fzd4 binds to Wnt11 in neonatal cardiomyocytes and in the mouse heart. Third, Fzd4 loss in neonatal cardiomyocytes stimulated neonatal cardiomyocyte proliferation and cell cycle activity mimicking the impact of Wnt11 loss. Fourth, Fzd4 loss suppressed the Wnt11-Rb1 axis by increasing p-Rb1. Last, exogenous rWnt11 treatment failed to mitigate the impact of Fzd4 loss by rescuing Rb1 activity. Together, the data integrates the Wnt11/Fzd4-Rb1 signaling axis with neonatal cardiomyocyte cell cycle exit and maturation mechanisms.

Neonatal heart maturation in mammals is a precisely controlled process in a chamber-specific, temporally regulated manner [1-4]. FZD proteins are 7-pass transmembrane proteins and members of the G protein-coupled receptor (GPCRs) superfamily [14]. Fzds are functionally classified as WNT receptors and show specific and dynamic expression during different embryonic development stages, playing essential roles in the regulation of developmental processes such as cell specification, cell polarity, neural patterning, and vascular development [14-17]. The dynamic regulation of Fzd4 expression in neonatal heart mirroring Wnt11 regulation and expression peak in RV at P3, supports its prioritization among the eight known Fzd receptors that have been characterized to date. Nevertheless, it has been shown that the Fzd proteins can be redundant at the functional level, and Wnt11 may signal via other Fzd receptors [15]. Contrary to this notion, the outcomes of Fzd4 loss in neonatal cardiomyocytes and the failure of rWnt11 to restore Rb1 activity support Fzd4 to be a specific, required, and non-redundant receptor for the Wnt11-Rb1 signaling axis in neonatal cardiomyocytes. Moreover, co-IP analysis confirmed the direct ligand-receptor relationship in neonatal cardiomyocytes *in vitro and in vivo*.

Our results show that Wnt11 and Fzd4 are both suppressed by perinatal hypoxia in an RV-specific manner [Figure 2], indicating that Wnt11 signaling via Fzd4 provides a coordinated link between the early postnatal environment and the cell cycle exit in the mammalian heart chambers, and identifies Wnt11/Fzd4-Rb1 as an important regulator of cardiomyocyte perinatal transition from proliferative to non-proliferative mature phenotype. The disruption of this regulatory axis under the influence of hypoxic environment indicates a reversion to a more immature proliferative state of cardiomyocytes. The resulting less mature proliferative phenotype of neonatal cardiomyocytes may have a negative impact by delaying neonatal heart maturation and contribute to the overall RV pathology in cyanotic CHDs, such as TOF and other forms of cyanotic CHDs [1,3,22]. On the other hand, it is tempting to propose that this outcome is potentially desired and a viable target for novel regenerative interventions [20,21] to enhance neonatal heart growth in the context of hypoplastic left ventricle (HLHS) and other forms of single ventricle physiology. Awaiting future confirmatory studies in mice and humans to affirm the pathological effect vs the therapeutic values, the findings from this study provide important insights into the signaling players of a novel gene-environment regulatory circuit in neonatal cardiomyocyte maturation and cell cycle exit in neonatal hearts that holds translational relevance for vulnerable newborn infants with CHDs [22].

## Methods

### Experimental animals

The experiments were conducted as part of an ongoing study under an active animal protocol approved by the Institutional Animal Care and Use Committee of UCLA. Animal handling followed the standards of the Guide for the Care and Use of Laboratory Animals. Pathogen-free male and female C57BL/6J mice from the inbred C57BL/6 strain were obtained from Charles River Laboratories. Successful mating was confirmed by plug detection. Timed pregnant dams were kept in cages under a 12-hour light/dark regime with food and water ad libitum. At E19.5, caesarian section delivery was performed, and the hearts from male pups were excised to achieve the P0 time point (as specified below). The remaining dams were allowed to deliver normally. Male pups and their mothers were kept in the same condition until the predetermined specific end point. Perinatal systemic hypoxia exposure (FIO2 10%) was performed following a previously established protocol [1]. The dams were maintained with their offspring until the predefined time point (P0, P3, or P7), whereas the dams carrying the control neonates (normoxia) were maintained in an ambient environment. Male pup hearts were excised at 3 time points of perinatal circulatory transition: P0 (birth before the ductal closure), P3 (establishing postnatal circulation), and P7 (terminal differentiation of most CMCs based on current literature evidence). RV and LV tissues were separated en bulk from each heart, leaving the septum out. The samples were snap frozen in liquid nitrogen and stored separately at –80°C.

### Gene expression analysis by qRT-PCR

Total RNA was isolated from pooled male pups’ LV and RV separately, using an RNeasy Mini Kit (QIAGEN) per manufacturer’s protocol. DNA contamination was eliminated using an RNase-Free DNase Set (QIAGEN). For reverse transcription, 1 μg of total RNA was used to generate first-strand cDNA with oligo-dT primers. Real-time PCR was performed using the SYBR Green Mix (Bio-Rad) on the CFX96 Real-time System (Bio-Rad). qRT-PCR was performed using housekeeping genes GAPDH for normalization and significant differences in gene expression were reported based on fold change. Primer sequences are listed in Supplemental Table 1.

### Immunohistochemistry

Cells were fixed in 4% (v/v) formaldehyde in PBS for 20 minutes. After rinsing with PBS, the cells were incubated with 0.1% Triton X-100 in PBS for 15 minutes at room temperature and rinsed. Heart tissue was fixed in 4% (v/v) formaldehyde, embedded in paraffin, and cut into 5-μm-thick tissue sections. After deparaffinization, slides were subjected to antigen retrieval. After blocking PBS containing 10% bovine serum albumin for 1 hour, the cells and tissue sections were incubated with primary antibodies overnight and then appropriate AlexaFluor-conjugated secondary antibodies (Life Technologies) for 1 hour. Cell nuclei were eventually counterstained with DAPI in the mounting medium (Life Technologies). Images were recorded on a Nikon confocal microscope. To count pH3– positive CMCs, we carefully examined the sections under confocal microscopy to ensure the counted pH3– positive nuclei were fully embedded in sarcomeres as expected for CMCs. Primary antibodies used to detect the expression of the proteins of interest are listed in Supplemental Table 2.

### Co-Immunoprecipitation Assays

Cells or mouse P3 hearts were lysed in RIPA buffer and the cell lysates were precleared with protein G– Dynabeads for 1 h at 4°C. Antibodies were conjugated to protein-G-Dynabeads for 10 min at room temperature. Proteins were then immunoprecipitated by incubating the cell lysates for 2 hours at 4°C with the anti-Wnt11 antibody and the Normal IgG conjugated protein-G-Danabeads. The presence of FZD4 in the precipitated material was assessed by western blotting with anti-FZD4 antibody.

### Western blot analysis

After protein quantification, protein samples were electrophoresed in 4%–12% SDS polyacrylamide gels before transfer to PVDF membranes (Millipore). HRP-conjugated secondary antibodies (Bio-Rad) were used, followed by ECL reaction to develop the blots according to the manufacturer’s instructions. Band intensities from film were analyzed by IMAGEQUANT 5.2 software (Molecular Dynamics) if needed. Primary antibodies used to detect the expression of the proteins of interest are listed in Table 2. NRVM culture, rWnt11, and Wnt11 siRNA treatment. NRVMs were isolated from 2-day-old rat pups and cultured in DMEM/5% (vol/vol) FBS with antibiotics at 37°C and 5% CO2. On the next day, cells were incubated with 200 ng/ml rWnt11 (R&D Systems) for 48 hours. For Wnt11 siRNA treatment, NRVMs were transfected in 6-well plates with 30 nM siRNA and 6 μl Lipofectamine RNAiMAX Transfection Reagent (Life Technologies) in Opti-MEM (Life Technologies). Culture medium was changed to DMEM/5% FBS after 4 hours. After 48 hours, total RNA was obtained using TRIzol reagent (Life Technologies), and total protein was isolated using cell lysis buffer (Life Technologies). For identification of phosphorylated proteins, Phosphatase Inhibitor Cocktail (Sigma-Aldrich) was added to the lysis buffer (Table 3).

### Statistics

Quantified results are presented as mean ± SEM. Student’s *t* test (unpaired, 2-tailed) and ANOVA with post-hoc Kruskal-Wallis were used for comparing 2 groups and more than 2 groups, respectively; *P* less than or equal to was considered significant, unless specified otherwise. The correlation of gene expression for each mRNA/trait pair was calculated using Pearson’s correlation and Benjamini-Hochberg correction methods. A Benjamini-Hochberg–adjusted correlation *P* value less than or equal to 0.05 was considered significant.

### Study Protocol Approval

All animal-related experimental protocols were approved by the UCLA Animal Care and Use Committee.

## Author contributions

MT conceived the project, designed and conducted the research, analyzed most of the data, and managed funding. XK performed experiments, generated figures, participated in experimental design, and wrote the manuscript draft. JM and CW participated in experiments, data analysis, and manuscript writing.

