## Supplemental Tables for "Wnt11 Positively Regulates Neonatal Cardiomyocyte Maturation at the Interphase of Life via Frizzled 4 Receptor"

| Supplemental Table 1. List of primers used for quantitative RT-PCR | | |
|---|---|---|
| Gene | Forward Primer | Reverse Primer |
| Fzd4 (Mouse, Rat) | | |
| Wnt11 (Mouse, Rat) | GCTGGCACTGTCCAAGACTC | CTCCCGTGTACCTCTCTCCA |
| AurkB (Mouse, Rat) | GGCTCAGAAAGAGAACGTCTACCCG | CAGTCAACTTCTGACCAGGGACAGC |

| Supplemental Table 2. List of antibodies and their sources | |
|---|---|
| Gene | Antibodies Source |
| Fzd4 | Abcam, ab83042 |
| Wnt11 | OriGene, TA308507 |
| Rb1 | Abcam, ab181616 |
| p-Rb1 (s795) | Cell Signaling |
